# Modeling the Correlation between *Z* and *B* in an X-ray Crystal Structure Refinement

**DOI:** 10.1101/2023.07.04.547724

**Authors:** Trixia M. Buscagan, Douglas C. Rees

## Abstract

We have examined how the refined *B*-factor changes as a function of *Z* (the atomic number of a scatterer) at the sulfur site of the [4Fe:4S] cluster of the nitrogenase iron protein by refinement. A simple model is developed that quantitatively captures the observed relationship between *Z* and *B*, based on a Gaussian electron density distribution with a constant electron density at the position of the scatterer. From this analysis, the fractional changes in *B* and *Z* are found to be similar. The utility of *B*-factor refinement to potentially distinguish atom types reflects the *Z* dependence of X-ray atomic scattering factors; the weaker dependence of electron atomic scattering factors on *Z* implies that distinctions between refined values of *B* in an electron scattering structure will be less sensitive to the atomic identity of a scatterer than for the case with X-ray-diffraction. This behavior provides an example of the complementary information that can be extracted from different types of scattering studies.

## Introduction

We recently reported a series of selenium-incorporated nitrogenase iron (Fe) protein crystal structures in which mixed occupancies of sulfur and selenium were observed at the chalcogenide sites of the [4Fe:4X] cluster (X = S, Se (Buscagan *et al*., 2022)). Occupancy refinements of these sites were correlated with shifts in the *B*-factor, reflecting the well-recognized correlation between occupancy and *B*-factor parameters that is most frequently encountered with solvents during protein structure refinement (Watenpaugh *et al*., 1978, Kundrot & Richards, 1987, Bhat, 1989, Jensen, 1990). By fixing the *B*-factors of the X atoms to match the *B*-factors of the Fe atoms, consistent with observations on other metalloprotein systems (Wittenborn *et al*., 2018, Jeoung *et al*., 2022), we refined occupancies at the mixed chalcogenide sites with minimal residual density in the *F*_obs_ -*F*_calc_ difference maps.

From that study, we became interested in characterizing the correlation between the atomic number, *Z* (as a proxy for occupancy), and the refined *B*-factor, to address two questions:

i. for a given change in *Z*, how much does *B* change?
ii. can a physical model be devised that captures this relationship?

To address the first question, the sulfide ligands in the [4Fe:4S] clusters of nitrogenase Fe protein PDB data sets 7TPW and 7TPY (with resolutions of 1.18 Å and 1.48 Å, respectively) were modeled with different elements in place of sulfur, and the *B*-factors refined. In these structures, the sulfurs in the cluster were fully occupied; i.e., no selenium was present. PHENIX (Liebschner *et al*., 2019), and COOT (Emsley *et al*., 2010) were used for this analysis, and the cluster coordinates were fixed so that only the *B*-factors were refined. Neutral scattering factors were used for the cluster atoms. The refined *B* values are provided in Appendix A. Over a *Z* range from 7 (N) to 34 (Se), the expected positive correlations between *Z* and refined *B* values are evident for both structures (Figure 1). A linear fit to this data for 8 ≤ *Z* ≤ 25 indicates that over this range, *B* varies approximately linearly with changes in *Z*; the fractional change in *B* relative to the fractional change in *Z* was found to be approximately 0.9 and 0.8 for the 7TPW and 7TPY data sets, respectively (Appendix A). Thus, the answer to the first question is that for this system, a 10% change in *Z* results in an approximately 8-9% change in *B*.

**Figure 1.**
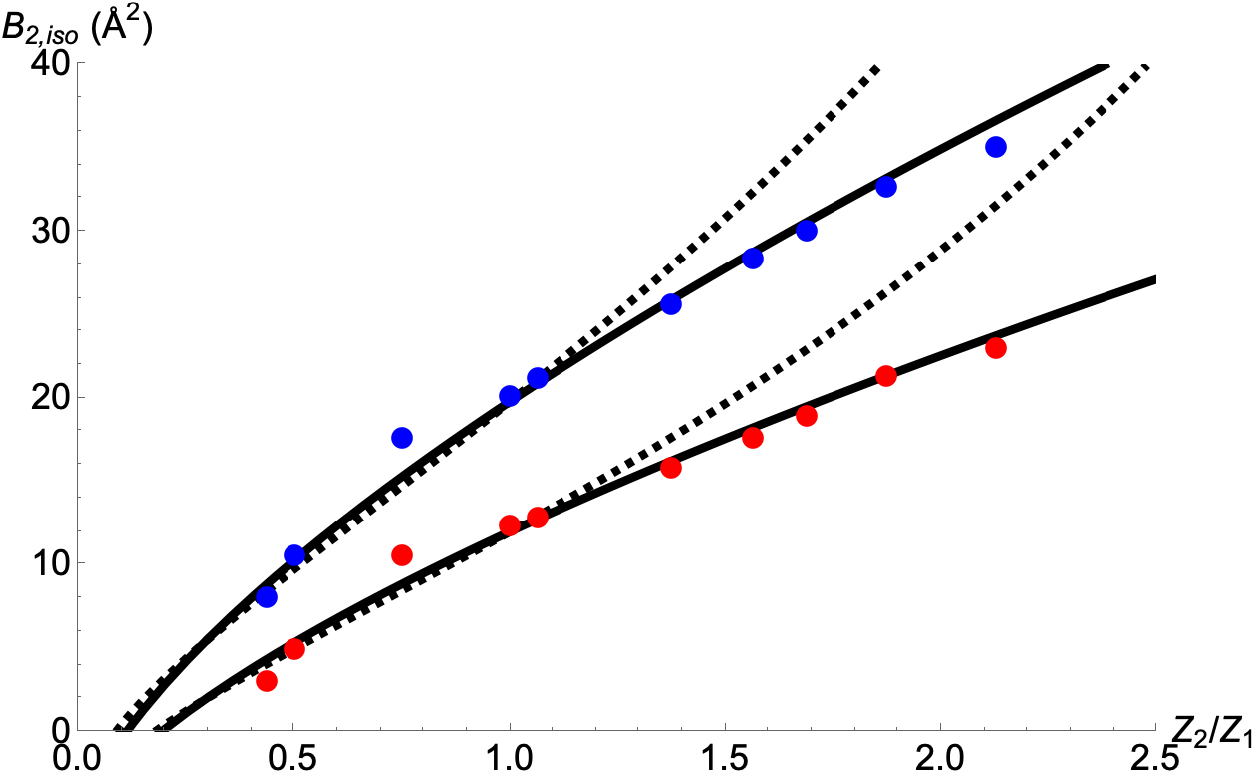
Refined *B*_2,iso_ values as a function of *Z*_2_/*Z*_1_, the ratio of the atomic number of the scatterer refined in the chalcogenide site (*Z*_2_) relative to *Z*_1_ = 16 (the true scatterer, sulfur), in the [4Fe:4S] cluster of the nitrogenase Fe protein (PDB data sets 7TPW (red circles) and 7TPY (blue circles)). The solid and dashed lines represent the fits to Eq. 4 and Eq. 6, respectively, with *Z*_1_ = 16 e^-^ and *B*_0_ = 6 Å^2^ for both structures, and *B*_1,iso_ = 12.0 Å^2^ and 19.8 Å^2^ for the 7TPW and 7TPY structures, respectively.

### Modeling the relationship between Z and B

To model the relationship between *Z* and *B*, we represent a scatterer by a single Gaussian with atomic number *Z* and overall temperature factor *B* (Teneyck, 1977). The electron density *ρ* (*r*) is then described:

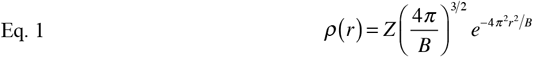

with

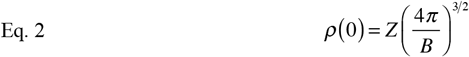

It is important to recognize that the *B* in Eq. 1 and Eq. 2 includes contributions from both the atomic scattering factor *B*_0_ and the isothermal temperature factor *B*_iso_, with *B* = *B*_0_ + *B*_iso_. From Eq. 2 and *ρ* (0) calculated with the Cromers and Mann atomic scattering factors (Cromer & Mann, 1968) and *B*_iso_ = 16 Å^2^, *B*_0_ is found to be approximately 8 Å^2^ and 6 Å^2^ for N and S, respectively. If the true *Z/B* for a given atom are *Z*_1_ and *B*_1_, but the refinement is conducted with *Z*_2_, the corresponding *B*_2_ will be shifted from the true value to compensate for the incorrect occupancy. We developed two simple models to capture the possible relationship between *Z*_*2*_ and *B*_*2*_:

**Model 1:** *B*_2_ is calculated for a given *Z*_2_ such that the density at the atomic position, *ρ* (0), has the same value as for *Z*_1_, *B*_1_. For a single Gaussian, this is equivalent to equating *ρ* (0), in Eq. 2 calculated for either *Z*_1_, *B*_1_ or *Z*_2_, *B*_2_, which gives

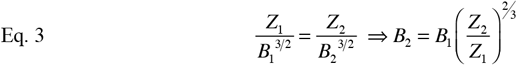

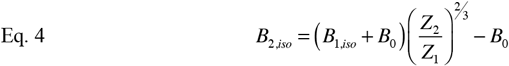

The ratio *Z*_2_*/Z*_1_ corresponds to the occupancy of the *Z*_1_ scatterer at the site (to within the approximation that the shape of the atomic scattering factor is independent of *Z*).

**Model 2:** In this case, *B*_2_ is calculated for a given *Z*_2_ to minimize the square of the difference density over the atomic volume:

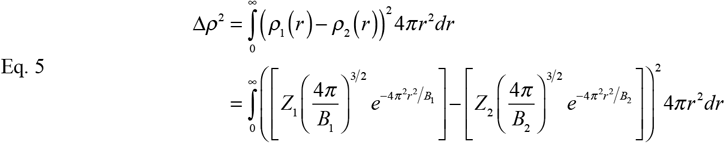

From the condition that 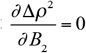 at the minimum, one can derive (Appendix B)

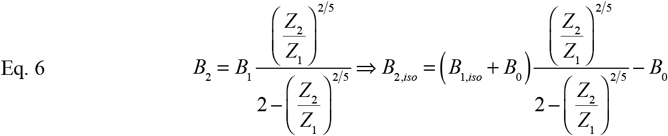

The variations in *B*_2,iso_ as a function of *Z*_2_/*Z*_1_ were evaluated from Eq. 4 and Eq. 6 (Figure 1). For these calculations, *Z*_1_ = 16 e^-^ and *B*_0_ = 6 Å^2^, with *B*_1,iso_ = 12.0 Å^2^ and 19.8 Å^2^ for the 7TPW and 7TPY structures, respectively. (These *B*_1,iso_ values correspond to the average *B*-factor for the two Fe sites in each structure; Appendix A.) As illustrated in Figure 1, while both Eq. 4 and Eq. 6 fit the refined *B* values reasonably well for *Z*_2_/*Z*_1_ < 1, the fit of Eq. 4 is superior over the entire range tested. This was a surprising result to us, as we anticipated that the Δρ_2_ model would better capture the structure refinement process; instead, the isolated atom approximation (reflected in the upper limit of r = ∝ in Eq. 5) for a macromolecular structure refinement is evidently less accurate relative to the localized treatment implicit in the derivation of Eq. 4.

## Discussion

We recognize that the approximations used to derive the relationship between *Z* and *B* in Eq. 4 are inferior to the results of a full structure refinement. Nevertheless, Eq. 4 can provide a useful starting point to evaluate the relationship between *Z* and *B* for scatterers involving either species of unknown atomic identity (such as the original analysis of the interstitial ligand in the nitrogenase FeMo-cofactor (Einsle *et al*., 2002), that prompted the initial development of model 2); partial occupancy (such as solvents (Watenpaugh *et al*., 1978)); mixed atomic composition (exchange reactions or disorder, as in (Spatzal *et al*., 2015, Buscagan *et al*., 2022)); or incorrect modeling of residues adopting distinct flipped orientations, such as the side chains of asparagine and glutamine residues where flipping interchanges N and O atoms (Word *et al*., 1999, Weichenberger & Sippl, 2006).

The utility of *B*-factor refinement to potentially distinguish atom types reflects the *Z* dependence of X-ray atomic scattering factors. An instructive comparison may be drawn to electron atomic scattering factors that depend on the Coulomb potential of the scatterer and have a less significant dependence on *Z*. In particular, *ρ*(0) for fully occupied C, N and O atoms calculated using electron scattering factors (parameterized in (Saha *et al*., 2021)) with a *B* = 16 Å^2^ vary by less than 2%, while the corresponding variation between the C and O atoms using X-ray scattering factors is approximately 50%. Consequently, the distinctions between the C, N, and O atoms in an electron scattering map are less evident than in X-ray scattering maps. This property of electron scattering was reflected in our recent structure determination of an antibiotic peptide by micro-electron diffraction, where the orientation of a histidine sidechain in a novel cross link could not be established from an analysis of the *B*-factors for the two distinct rotamers (Miller *et al*., 2022). A “multi-messenger” approach using combinations of X-ray (with anomalous scattering, if applicable), electron, and neutron diffraction can provide additional experimental restraints to help resolve ambiguities arising in structure refinements from the correlation between *Z* and/or occupancy with the *B*-factor of scatterers.

## Acknowledgments

We thank Jens Kaiser for enlightening discussions. The authors are grateful to the Gordon and Betty Moore Foundation, Don and Judy Voet, and the Beckman Institute at Caltech for their generous support of the Molecular Observatory at Caltech. Use of the Stanford Synchrotron Radiation Lightsource, SLAC National Accelerator Laboratory, is supported by the U.S. Department of Energy, Office of Science, Office of Basic Energy Sciences under Contract No. DE-AC02-76SF00515. The SSRL Structural Molecular Biology Program is supported by the DOE Office of Biological and Environmental Research, and by the National Institutes of Health, National Institute of General Medical Sciences (including P41GM103393). This work was supported by the National Institute of Health (NIH Grant GM45162) and the Howard Hughes Medical Institute (HHMI).

## Appendix A: *B*-factor refinement of different scatterers in the sulfur sites of a [4Fe:4S] cluster

| 7TPW refinement |  |  |  |  |  |  |  |  |  |  |
| --- | --- | --- | --- | --- | --- | --- | --- | --- | --- | --- |
| X = element | N | O | Mg | S | Cl | Ti | Mn | Co | Zn | Se |
| atomic number (*Z*) | 7 | 8 | 12 | 16 | 17 | 22 | 25 | 27 | 30 | 34 |
| $Z_2/Z_1$ | 0.44 | 0.50 | 0.75 | 1.00 | 1.06 | 1.38 | 1.56 | 1.69 | 1.88 | 2.13 |
| cluster atom | refined *B* values (Å<sup>2</sup>) |  |  |  |  |  |  |  |  |  |
| Fe1 (Å<sup>2</sup>) | 11.82 | 11.90 | 11.99 | 12.02 | 12.11 | 12.07 | 12.12 | 12.28 | 12.27 | 12.30 |
| Fe2 (Å<sup>2</sup>) | 11.86 | 11.88 | 12.01 | 11.98 | 12.03 | 12.06 | 12.03 | 12.14 | 12.29 | 12.18 |
| X3 (Å<sup>2</sup>) | 3.41 | 5.38 | 10.85 | 12.75 | 13.30 | 16.23 | 18.18 | 19.42 | 21.87 | 23.77 |
| X4 (Å<sup>2</sup>) | 2.80 | 4.61 | 10.22 | 11.99 | 12.49 | 15.41 | 17.18 | 18.53 | 20.84 | 22.21 |
| X *B*<sub>avg</sub> (Å<sup>2</sup>) | 3.11 | 5.00 | 10.54 | 12.37 | 12.90 | 15.82 | 17.68 | 18.98 | 21.36 | 22.99 |
| *R*<sub>work</sub> | 14.20 | 14.16 | 14.11 | 14.07 | 14.09 | 14.12 | 14.14 | 14.17 | 14.24 | 14.24 |
| *R*<sub>free</sub> | 16.09 | 16.03 | 15.92 | 15.89 | 15.84 | 16.06 | 16.04 | 16.04 | 16.34 | 16.34 |

| 7TPY refinement |  |  |  |  |  |  |  |  |  |  |
| --- | --- | --- | --- | --- | --- | --- | --- | --- | --- | --- |
| X = element | N | O | Mg | S | Cl | Ti | Mn | Co | Zn | Se |
| cluster atom | refined *B* values (Å<sup>2</sup>) |  |  |  |  |  |  |  |  |  |
| Fe1 (Å<sup>2</sup>) | 19.82 | 19.73 | 19.91 | 19.73 | 19.91 | 19.77 | 19.60 | 19.60 | 19.44 | 19.38 |
| Fe2 (Å<sup>2</sup>) | 19.84 | 20.01 | 20.13 | 19.83 | 20.02 | 20.02 | 19.75 | 19.75 | 19.76 | 19.59 |
| X3 (Å<sup>2</sup>) | 8.33 | 10.86 | 17.83 | 20.38 | 21.47 | 26.02 | 28.68 | 30.40 | 33.01 | 35.31 |
| X4 (Å<sup>2</sup>) | 7.90 | 10.39 | 17.38 | 19.81 | 20.96 | 25.40 | 28.03 | 29.80 | 32.30 | 34.76 |
| X *B*<sub>avg</sub> (Å<sup>2</sup>) | 8.12 | 10.63 | 17.61 | 20.10 | 21.22 | 25.71 | 28.36 | 30.10 | 32.66 | 35.04 |
| *R*<sub>work</sub> | 17.34 | 17.25 | 17.18 | 17.17 | 17.23 | 17.24 | 17.26 | 17.23 | 17.27 | 17.36 |
| *R*<sub>free</sub> | 19.21 | 18.98 | 18.85 | 19.01 | 18.92 | 19.02 | 19.03 | 19.00 | 19.15 | 19.06 |

The *B*-factors are refined as a function of the element in the chalcogenide site of the [4Fe:4S] cluster for two nitrogenase Fe protein structures (PDB IDs 7TPW and 7TPY, with resolutions of 1.18 Å and 1.48 Å, respectively)) detailed in (Buscagan *et al*., 2022). The cluster sits on a crystallographic two-fold axis, so that the crystallographically unique sites are designated Fe1, Fe2, X3 and X4. Least squares lines fit to the *B*_avg_ values of the scatterers at the X position over the range 8 ≤ *Z* ≤ 25 yielded slopes of 0.692 and 0.987 for the 7TPW and 7TPY data sets, respectively; when normalized to the appropriate *Z* and *B* values for S, the fractional changes in *B* with a fractional change in *Z* are found to be 0.896 and 0.786, respectively.

## Appendix B

Derivation of Equation 6

Eq. 6 can be derived from Eq. 5 as follows. The integral and derivative were evaluated with Mathematica^®^.

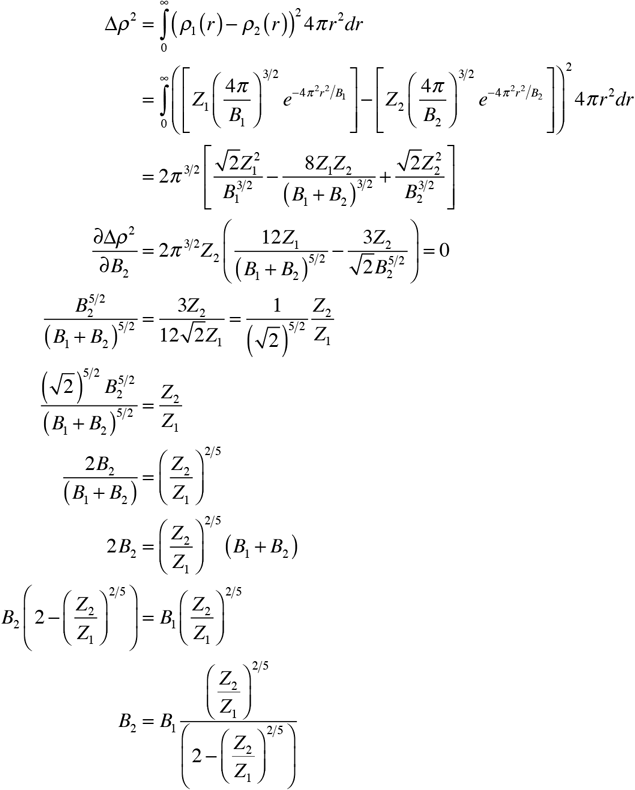

